# The mitochondrial dicarboxylate carrier mediates in vivo hepatic gluconeogenesis

**DOI:** 10.1101/2024.09.12.612761

**Authors:** Daniel J. Pape, Kelly C. Falls-Hubert, Ronald A. Merrill, Adnan Ahmed, Qingwen Qian, Gavin R. McGivney, Paulina Sobieralski, Adam J. Rauckhorst, Ling Yang, Eric B. Taylor

**Author notes:** Authors contributed equally (Daniel J. Pape, Kelly C. Falls-Hubert). Correspondence (E.B.T).

## Abstract

Hepatic gluconeogenesis (GNG) is essential for maintaining euglycemia during prolonged fasting. However, GNG becomes pathologically elevated and drives chronic hyperglycemia in type 2 diabetes (T2D). Lactate/pyruvate is a major GNG substrate known to be imported into mitochondria for GNG. Yet, the subsequent mitochondrial carbon export mechanisms required to supply the extra-mitochondrial canonical GNG pathway have not been genetically delineated. Here, we evaluated the role of the mitochondrial dicarboxylate carrier (DiC) in mediating GNG from lactate/pyruvate. We generated liver-specific DiC knockout (DiC LivKO) mice. During lactate/pyruvate tolerance tests, DiC LivKO decreased plasma glucose excursion and ^13^C-lactate/-pyruvate flux into hepatic and plasma glucose. In a Western diet (WD) feeding model of T2D, acute DiC LivKO after induction of obesity decreased lactate/pyruvate-driven GNG, hyperglycemia, and hyperinsulinemia. Our results show that mitochondrial carbon export through the DiC mediates GNG and that the DiC contributes to impaired glucose homeostasis in a mouse model of T2D.

## INTRODUCTION

Gluconeogenesis (GNG), the de novo synthesis of glucose, is essential for maintaining euglycemia during prolonged fasting. The liver is the primary gluconeogenic organ, with minor contributions from the kidney and small intestine ^1,2^. In T2D, hepatic insulin resistance leads to chronically elevated GNG that drives hyperglycemia contributing to cardiovascular disease and other co-morbidities that increase mortality ^3–10^. Furthermore, current treatments for T2D do not adequately restrain GNG, as evidenced by a correlation between gluconeogenic rates and fasting blood glucose concentrations in T2D patients receiving standard-of-care treatment ^11–13^. Thus, improved methods to modulate GNG are needed for the effective treatment of T2D. However, a basic limiter for the design of new therapeutic strategies is an incomplete understanding of the metabolic pathways and mechanisms supplying the carbon building blocks for GNG.

Circulating lactate/pyruvate is a major substrate for hepatic GNG ^14,15^. The liver takes up lactate/pyruvate produced by systemic metabolism, where it is oxidized to pyruvate by cytosolic lactate dehydrogenase. Mitochondria then import pyruvate through the mitochondrial pyruvate carrier (MPC), where it may be converted to oxaloacetate (OAA) by pyruvate carboxylase (PC) for anaplerotic GNG flux or into acetyl-CoA by pyruvate dehydrogenase (PDH) for forward TCA cycle flux. During fasting, hepatic PC activity greatly exceeds PDH, which favors pyruvate entry into GNG ^16–18^. In T2D, carbon flux through the MPC and PC increases, facilitating excessive GNG ^19,20^. Nevertheless, because the reactions of the central GNG pathway occur outside mitochondria, pyruvate-derived carbon must be exported for continued flux into GNG. However, the mitochondrial carbon export mechanisms supplying GNG have not been genetically defined.

Mitochondrial malate export through the mitochondrial dicarboxylate carrier (DiC), now known to be encoded by the *SLC25A10* gene, has been hypothesized to link mitochondrial anaplerosis with GNG ^16,21^. By this model, pyruvate is imported into mitochondria through the MPC and anaplerotically channeled into malate through PC and malate dehydrogenase (MDH). Malate is then transported out of mitochondria in exchange for inorganic phosphate, converted to oxaloacetate, and committed to GNG after decarboxylation and phosphorylation to phosphoenolpyruvate by phosphoenolpyruvate carboxykinase (PEPCK). Mitochondrial malate is biochemically well-suited to function as a GNG precursor because of its relatively large pool size, sequestration away from citrate synthesis, and export through the DiC by exchange with inorganic phosphate. Notably, the latter makes the DiC unique among known mitochondrial carriers in being able to export four net carbons ^22,23^. Given that TCA cycle anaplerotic fluxes and GNG both increase in T2D, the DiC is well-positioned to contribute to the excessive GNG of T2D ^19^.

To understand the role of the DiC in hepatic GNG, we generated liver-specific DiC knockout mice. Using both substrate tolerance tests and ^13^C stable-isotope tracing, we show that hepatic DiC deletion impairs glucose production from lactate/pyruvate. Furthermore, we demonstrate that hepatic DiC deletion in the Western diet feeding model of T2D mitigates hyperglycemia and hyperinsulinemia. Overall, our results show that the hepatic DiC mediates GNG in vivo and contributes to the pathophysiology of T2D.

## RESULTS

### Whole-body DiC knockout-first (KO1) mice have decreased body weight at weaning but grow normally

The overarching hypothesis of this investigation is that the DiC mediates hepatic GNG from pyruvate (Figure 1A). To understand both the physiologic context for and the role of the hepatic DiC in GNG, we generated mice with splice-disrupted, “knockout first” (KO1) *tm1a* (*DiC^KO1^*) and conditional, floxed *tm1c* (*DiC^fl^*) *Slc25a10* alleles from cryopreserved sperm obtained from EUCOMM (Figure 1B) ^24^. To obtain whole-body DiC KO1 mice, *DiC^+/KO1^* mice were intercrossed. *DiC ^KO1/KO1^* mice were viable, though body weight at weaning was significantly decreased in both males and females (Fig 1C-D). Nonetheless, body weight was normalized by 6 weeks of age, suggesting that whole-body DiC insufficiency may impair development but is not detrimental to mouse growth and survival.

**Figure 1:**
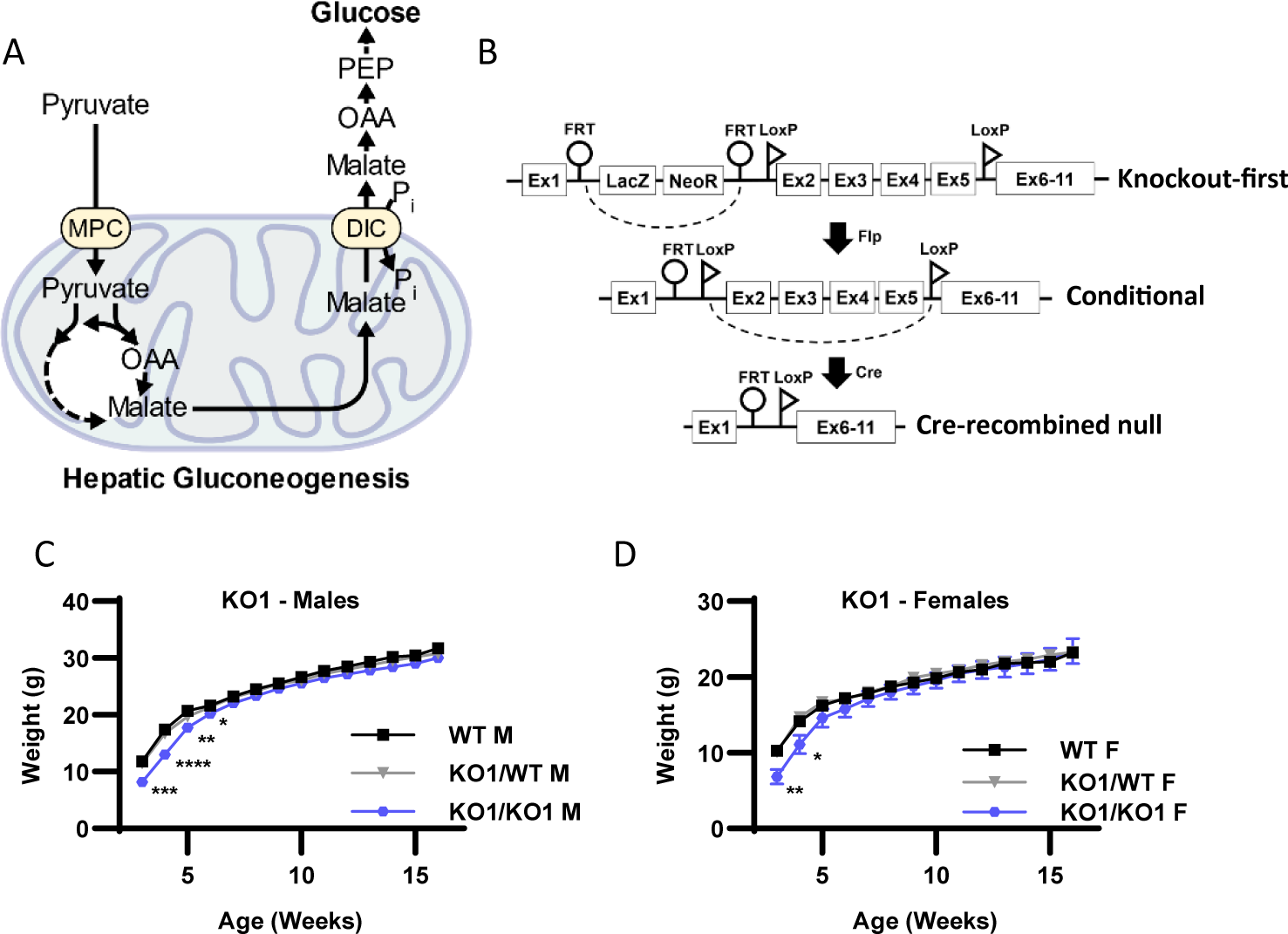
Whole-body DiC knockout-first (KO1) mice have decreased body weight at weaning but grow normally. A. Schematic illustrating the role of the mitochondrial dicarboxylate carrier (DiC) in gluconeogenesis. MPC, mitochondrial pyruvate carrier; OAA, oxaloacetate; PEP, phosphoenolpyruvate. B. Schematic illustrating *Slc25a10^Tm1a^* knockout-first allele (KO1), the conditional (*DiC^fl/fl^*) allele, and the AAV-TBG-Cre mediated, liver-specific null allele (DiC LivKO). C-D. Body weight of wild type (WT), heterozygous (KO1/WT) and knockout-first (KO1/KO1) male (C) and female (D) mice, beginning on the day of weaning (3 weeks of age). Data are presented as mean ± SEM. Statistical significance evaluated by one-way ANOVA within each time point denoted as * p<0.05, ** p<0.01, *** p<0.001, **** p<0.0001. N for male mice = 11-18 WT, 26-39 KO1/WT, and 13-15 KO1/KO1. N for female mice = 13-17 WT, 15-34 KO1/WT, and 8-9 KO1/KO1.

### DiC LivKO decreases glucose excursion during lactate/pyruvate tolerance tests

To selectively test the role of the DiC in hepatic GNG, we generated liver-specific DiC knockout (DiC LivKO) and control (WT) mice by injecting *DiC^fl/fl^* mice with AAV-TBG-Cre or AAV-TBG-NULL (EV). Western blots of isolated mitochondria from liver and kidney tissue obtained 10 weeks post-injection showed liver-selective DiC knockout (Fig 2A). DiC LivKO did not significantly alter body composition as measured by NMR (Fig 2B). Given our hypothesis that the DiC contributes to glucose homeostasis by mediating GNG, we performed metabolic tolerance tests by intraperitoneal injection, dosed by body weight. DiC LivKO did not alter glucose (1.25 g/kg) or insulin (0.75 U/kg) tolerance in healthy, normal chow (NCD) fed mice (Fig 2C-D). However, lactate/pyruvate (10:1 ratio to approximate endogenous redox balance; 2.2 g/kg or 3 g/kg lean mass) tolerance tests (L/PTT) showed that DiC LivKO decreased lactate/pyruvate-driven glucose excursion in vivo in both male and female mice (Fig 2 E-H).

**Figure 2:**
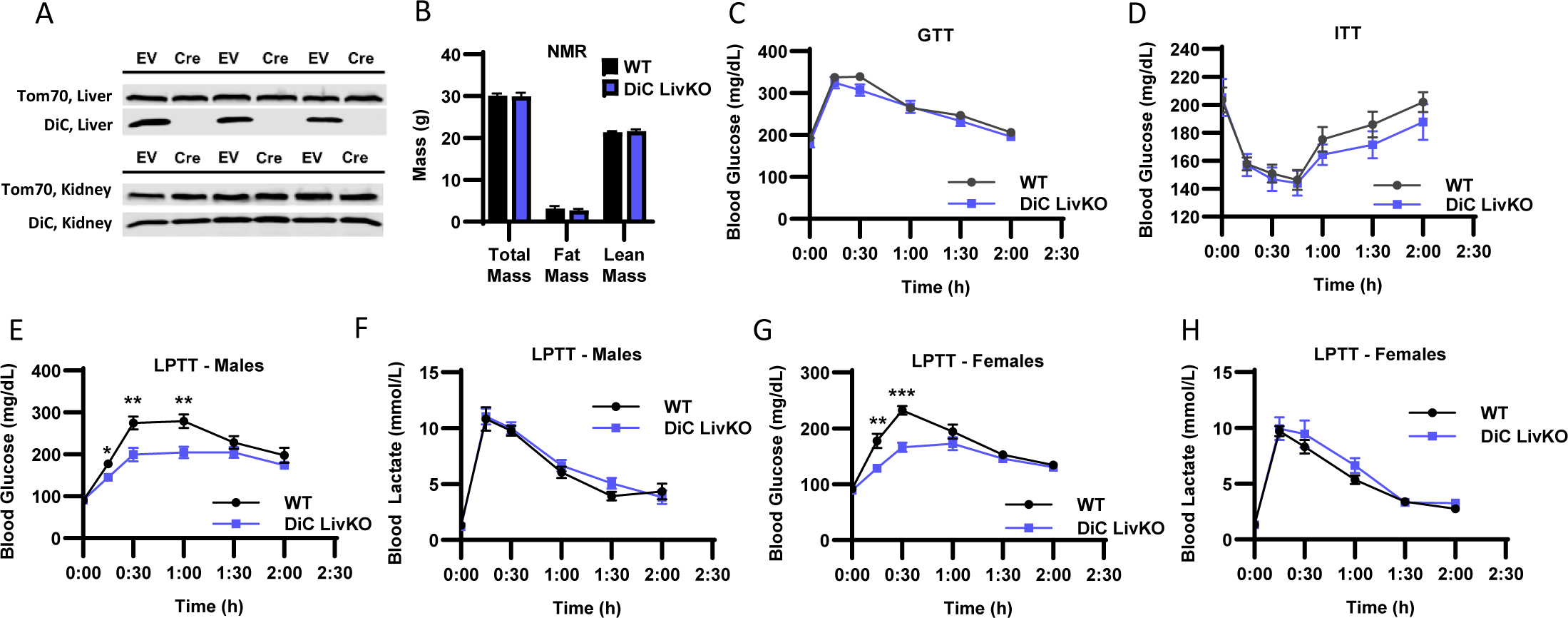
DiC LivKO decreases glucose excursion during lactate/pyruvate tolerance tests. A. Western blot of isolated liver and kidney mitochondria from AAV-TBG-NULL (WT) or AAV-TBG-Cre (DiC LivKO) injected *DiC^fl/fl^*mice, 10 weeks post-injection. B. Total body, fat, and lean mass determined by Nuclear Magnetic Resonance (NMR) of 17-18 week old WT and DiC LivKO mice, 9 weeks post-AAV EV/Cre injection. C. Blood glucose measured before and during an Intraperitoneal Glucose Tolerance Test (GTT, 1.25 g/kg body weight) performed on 11-12 week old WT and DiC LivKO mice, 3 weeks post-AAV EV/Cre injection. D. Blood glucose measured before and during an Insulin Tolerance Test (ITT, 0.75 U/kg body weight) performed on 12-13 week old WT and DiC LivKO mice, 4 weeks post-AAV EV/Cre injection. E-F. Blood glucose (E) and lactate (F) measured before and during a Lactate/Pyruvate Tolerance Test (L/PTT, 3.0 g/kg lean mass, 10:1 lactate/pyruvate) performed on 18-19 week old male WT and DiC LivKO mice, 10 weeks post-AAV EV/Cre injection. G-H. Blood glucose (G) and lactate (H) measured before and during a L/PTT (2.2 g/kg body weight, 10:1 lactate/pyruvate) performed on 13-14 week old female WT and DiC LivKO mice, 4 weeks post-AAV EV/Cre injection. Data are presented as mean ± SEM. Statistical significance evaluated by t-test at each time point denoted as * p<0.05, ** p<0.01, is *** p<0.001. N = 10 per group.

### Complementing DiC LivKO with ectopically expressed WT but not transport null R261Q DiC rescues glucose excursion during lactate/pyruvate tolerance tests

To test whether the effects of DiC LivKO on GNG are specific to DiC transport function versus a potential protein scaffolding role, we performed KO rescue experiments in vivo. Wild type DiC (DiC WT) and DiC R261Q point mutant constructs (DiC R261Q) were ectopically expressed in the liver by AAV8 administration and driven by the TBG promoter. Based on the orthologous R275Q mutation in the mitochondrial ornithine carrier (Orc1 encoded by the *Slc25a15* gene), we hypothesized that an R261Q mutation of DiC would ablate DiC transport function. The Orc1 R275Q mutation disrupts transport function without decreasing protein abundance or impairing mitochondrial localization ^25–27^. Additionally, the DiC AlphaFold model predicts that R261 resides in the central pore (Fig 3A), consistent with a role in transport function ^28,29^. Western blots of isolated liver mitochondria confirmed ectopic expression of DiC WT and R261Q at approximately endogenous WT levels (Fig 3B).

**Figure 3:**
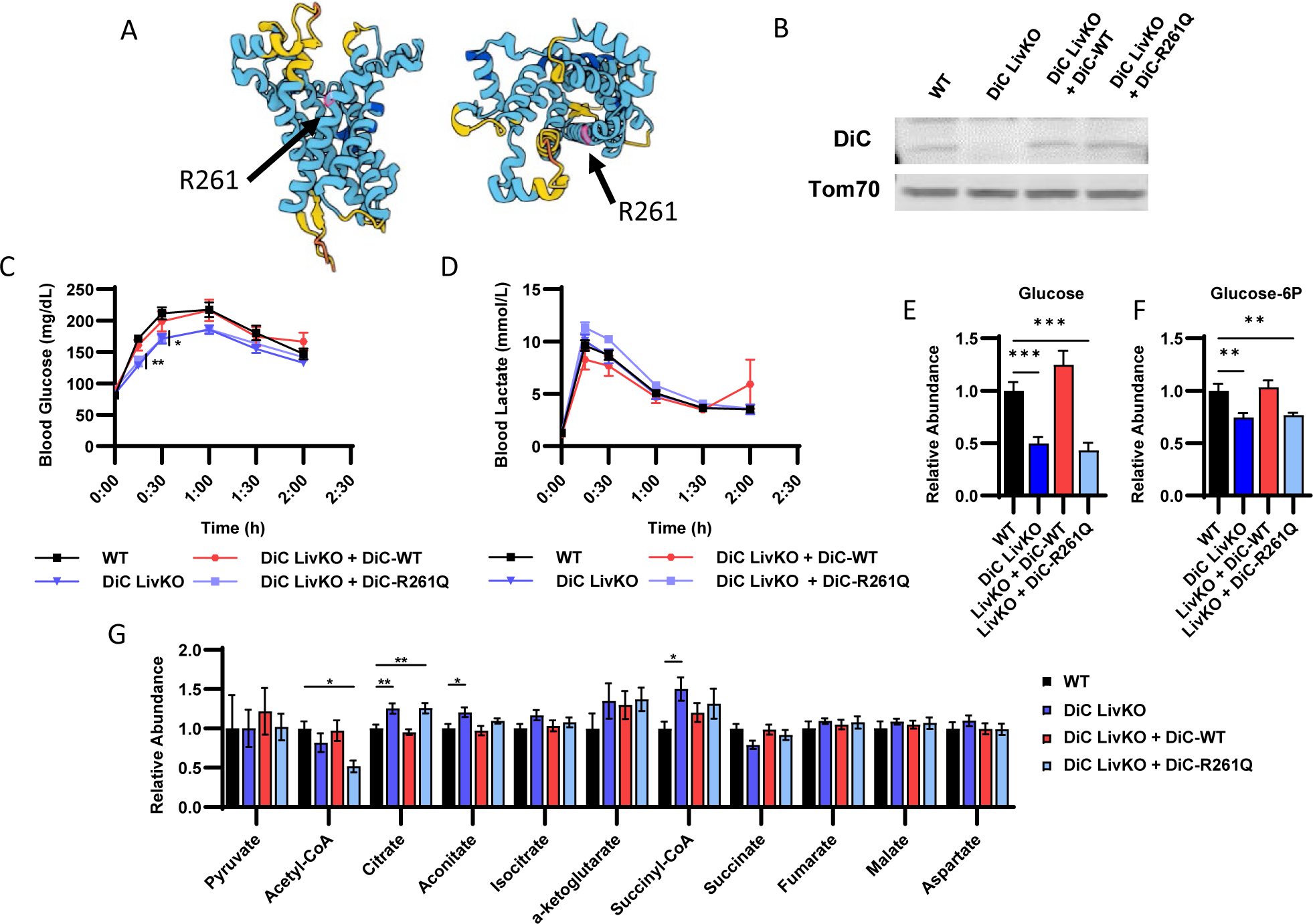
Complementing DiC LivKO with ectopically expressed WT but not transport null R261Q DiC rescues glucose excursion during lactate/pyruvate tolerance tests. A. AlphaFold structure of DiC protein denoting the expected location of R261 in the pore region. B. Western blot of isolated liver mitochondria from WT, DiC LivKO, DiC LivKO + DiC-WT, or DiC LivKO + DiC-R261Q mice. C-D. Blood glucose (C) and lactate (D) measured before and during a Lactate/Pyruvate Tolerance Test (L/PTT, 2.2 g/kg body weight, 10:1 lactate/pyruvate) performed on 10-11 week old WT, DiC LivKO, DiC LivKO + DiC-WT, or DiC LivKO + DiC-R261Q mice, 3 weeks post-AAV EV/Cre injection. E-G. Liver glucose (E), glucose-6P (F), and TCA cycle intermediates (G) levels were measured by LC-MS metabolomic profiling of 12-13 week old WT, DiC LivKO, DiC LivKO + DiC-WT, or DiC LivKO + DiC-R261Q mice, 6 weeks post-AAV EV/Cre injection. Data are presented as mean ± SEM. Statistical testing by one-way ANOVA and the Holm-Sidak post-hoc multiple comparison test within each time point compared to WT (L/PTT) and for each metabolite (LC-MS metabolomics). Significance denoted by * p<0.05, ** p<0.01, *** p<0.001. N = 10 per group.

To assess how complementation of DiC LivKO livers with DiC WT and DiC R261Q affects GNG, we performed lactate/pyruvate (10:1 lactate/pyruvate; 2.2 g/kg) tolerance tests (LPTTs). DiC LivKO again significantly decreased glucose excursion, which was reversed by DiC WT (Fig 3C-D). Conversely, DiC R261Q did not restore lactate/pyruvate-driven GNG (Fig 3C-D). These results suggest that the DiC transport activity is critical for GNG.

To test the effects of DiC LivKO plus complementation with DiC WT or DiC R261Q on the steady-state, post-absorptive liver metabolome, we performed metabolomic analysis of liver tissue collected from 4h-fasted mice. Notably, DiC LivKO decreased glucose and glucose-6P abundance, which was reversed by DiC WT but not R261Q (Fig 3E-F, Table S1). DiC LivKO also increased citrate and decreased acetyl-CoA abundances, consistent with oxaloacetate being directed away from reduction to malate and instead toward citrate synthesis by condensation with acetyl-CoA (Fig 3G, Table S1).

### DiC LivKO decreases GNG from ^13^C-lactate/-pyruvate

To more specifically test the role of the DiC in lactate/pyruvate-driven GNG, we employed stable isotope tracing with uniformly labeled (U) ^13^C-lactate/-pyruvate (9:1 ratio, 50% U^13^C-labeled, 2.2 g/kg). At 30 minutes post intraperitoneal injection, DiC LivKO mice exhibited decreased blood glucose and elevated blood lactate levels (Fig 4A-B, Table S2). Abundances of lactate and pyruvate in liver tissue were not significantly affected by DiC LivKO, though lactate percent enrichment was minimally lower, possibly due to tracer dilution from impaired clearance of endogenous lactate (Fig 4 C-D, Table S2). Similar to blood glucose levels, DiC LivKO also decreased abundance of hepatic glucose and glucose-6P as well as their percent M+2 and M+3 isotopologues, signifying decreased U^13^C-lactate/-pyruvate channeling into GNG (Fig 4 E-H, Table S2). In contrast, DiC LivKO led to increased abundance and percent enrichments of TCA cycle metabolites (Fig 4I-J, Table S2). These findings suggest that ^13^C-pyruvate is adequately imported into mitochondria for TCA anaplerosis and forward flux but that downstream GNG precursor(s) are not adequately exported.

**Figure 4:**
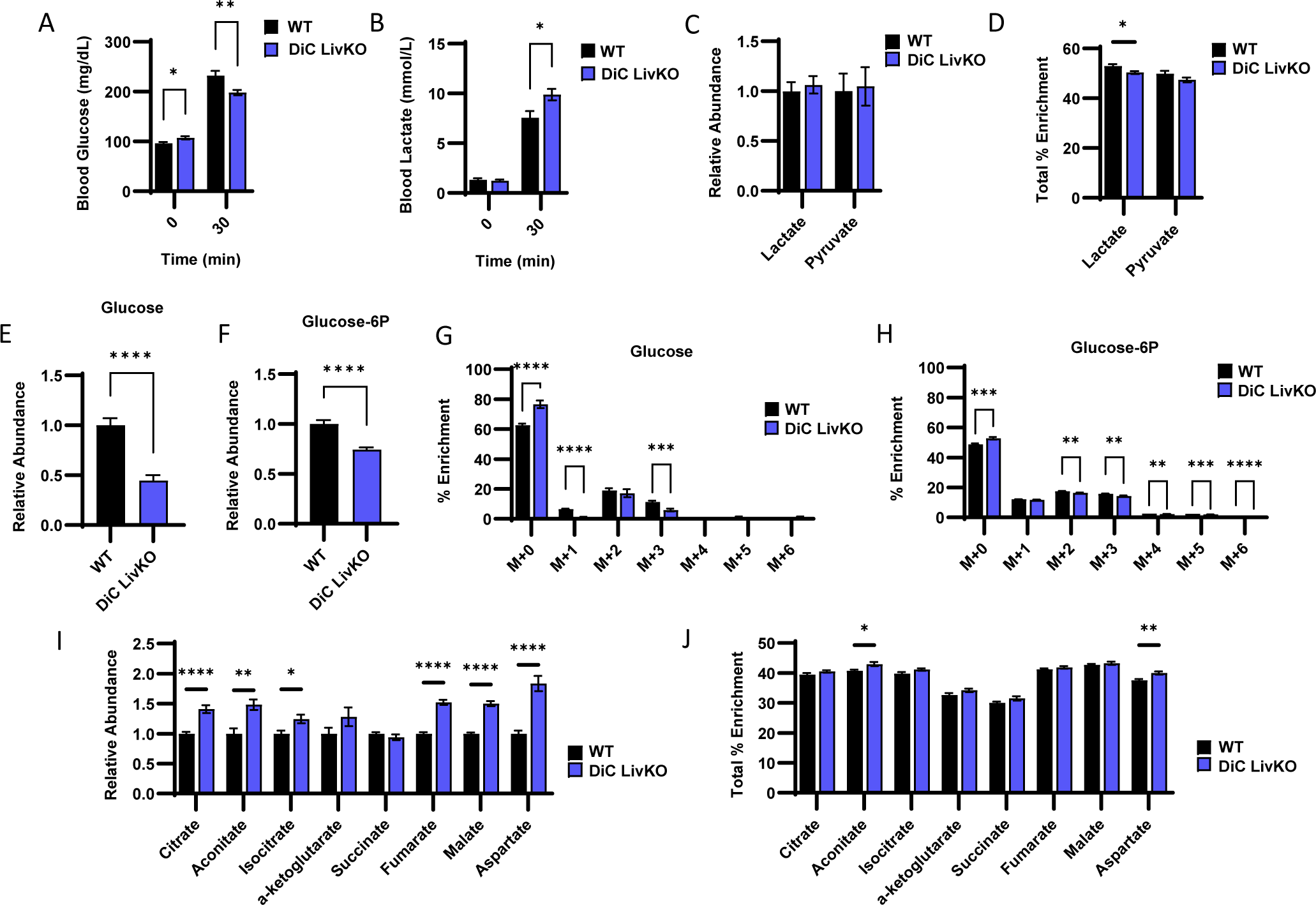
DiC LivKO decreases gluconeogenesis from ^13^C-lactate/-pyruvate. 13-14 week old WT and DiC LivKO mice were administered ^13^C-lactate/pyruvate tracer (2.2 g/kg body weight, 9:1 lactate/pyruvate) and euthanized 30 minutes later, 5 weeks post-AAV EV/Cre injection. A-B. Blood glucose (A) and lactate (B) measured by strip meter before tracer administration and at euthanasia. C-D. Terminal liver lactate and pyruvate levels (C) determined as the sum of all isotopologue peak areas and total percent enrichment (D) determined as the sum of all ^13^C labeled isotopologues) measured by LC-MS. E-H. Terminal liver glucose (E and G) and glucose-6P (F and H) levels (E and F) and mass isotopologue distribution analysis (G and H) measured by LC-MS. I-J. Terminal liver TCA cycle intermediate levels (I) and total percent enrichment (J) measured by LC-MS. Data are presented as mean ± SEM. Statistical analysis by t-test for each time point (blood glucose and lactate), metabolite (LC-MS analysis), or isotopologue (MID analysis). Significance denoted as * p<0.05, ** p<0.01, *** p<0.001, **** p<0.0001. N = 10 for each group.

### DiC LivKO normalizes blood glucose and insulin levels in Western Diet (WD) treated mice

GNG is known to contribute to elevated blood glucose in type 2 diabetes (T2D). Thus, we considered whether acute DiC LivKO would alter glucose homeostasis in a high fat-, high sucrose-diet feeding model of T2D. First, before administration of AAV-Cre to delete the DiC, mice were fed a normal chow diet (NCD) or WD for 14 weeks to induce insulin resistance and glucose intolerance (Fig 5A, Table S3 containing p-values for all two-way ANOVA comparisons). During weeks 14-16 of NCD vs WD feeding, we assessed mice for differences in body composition and glucose homeostasis. As expected, compared to NCD mice, WD mice had elevated total and fat mass (Fig 5B). WD also led to increased post-absorptive and fasted blood glucose, insulin, and Homeostatic Model Assessment for Insulin Resistance (HOMA-IR = [Blood Glucose (mmol/L) * Plasma Insulin (µIU/mL) / 22.5]) scores (Fig 5C-E). WD did not lead to elevated lactate/pyruvate-driven GNG as reflected by glucose excursion during lactate/pyruvate tolerance tests (10:1 ratio, 3 g/kg lean mass). However, WD did increase blood lactate levels, which could lead to elevated lactate/pyruvate-driven GNG beyond the time-course of the test in WD mice (Fig 5F-G).

**Figure 5:**
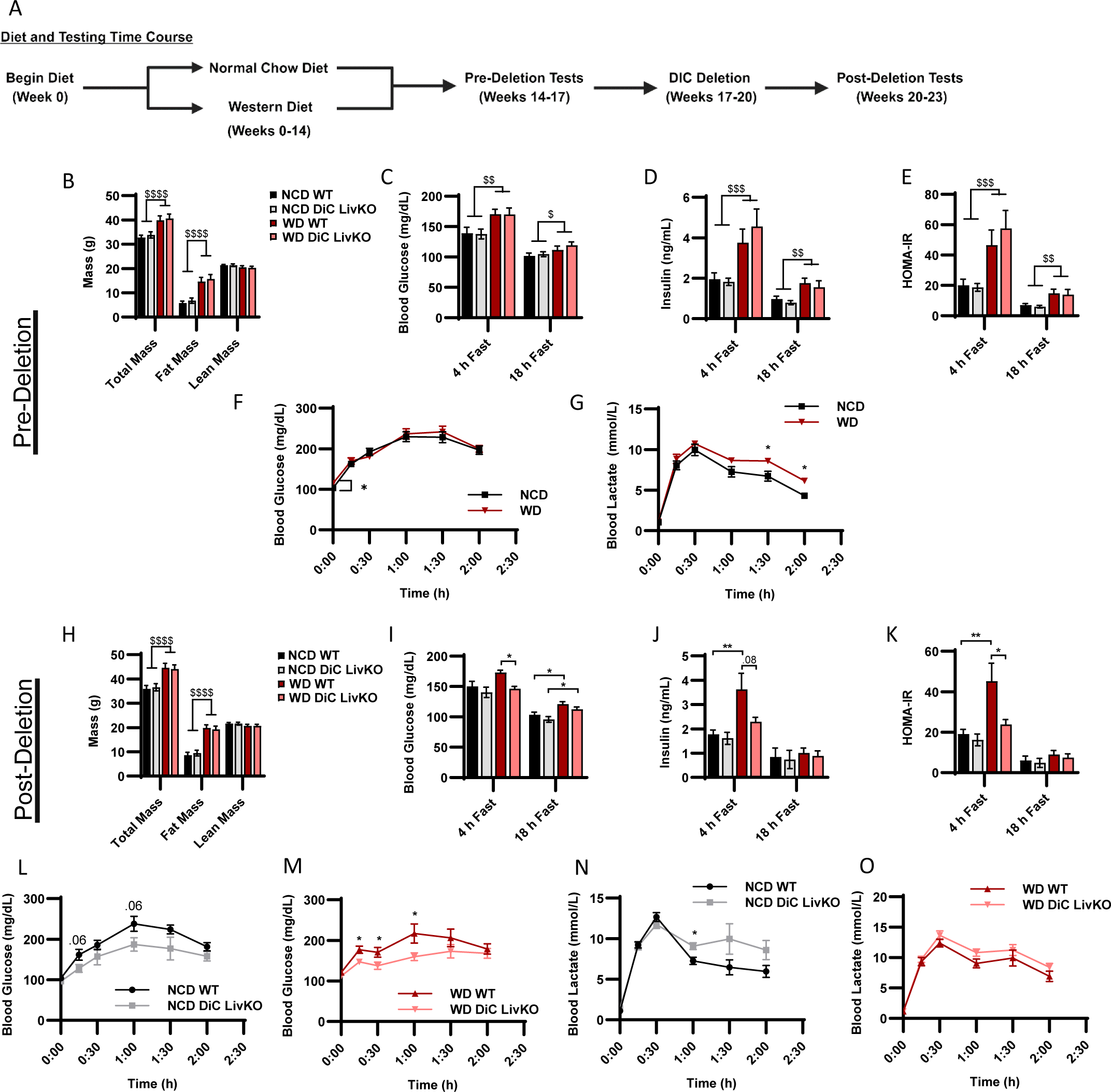
DiC LivKO normalizes blood glucose and insulin levels in Western Diet (WD) treated mice. A. Schematic illustrating the diet treatment and testing time course for *DiC^fl/fl^* mice. B. Total body, fat, and lean mass determined by Nuclear Magnetic Resonance (NMR) of 20-21 week old *DiC^fl/fl^* mice, fed normal chow (NCD) or western diet (WD) for 14 weeks. C-E. Blood glucose (C) and plasma insulin (D) measured in 4- and 18-hour fasted, 20-22 week old *DiC^fl/fl^*mice, fed NCD or WD for 14-15 weeks. The Homeostatic Model Assessment for Insulin Resistance (HOMA-IR, E) calculated from values shown in C and D. F-G. Blood glucose (F) and lactate (G) measured before and during a Lactate/Pyruvate Tolerance Test (L/PTT, 3.0 g/kg lean mass, 10:1 lactate/pyruvate) performed on 21-22 week old *DiC^fl/fl^* mice, fed NCD or WD for 15 weeks. H. Total body, fat, and lean mass determined by NMR of 26-27 week old WT and DiC LivKO mice, 3 weeks post-AAV EV/Cre injection, fed NCD or WD for 20 weeks. I-K. Blood glucose (I) and plasma insulin (J) measured in 4- and 18-hour fasted, 26-28 week old WT and DiC LivKO mice, 3-4 weeks post-AAV EV/Cre injection, fed NCD or WD for 20-21 weeks. HOMA-IR (K) calculated from values shown in I and J. L-O. Blood glucose (L and M) and lactate (N and O) measured before and during a Lactate/Pyruvate Tolerance Test (L/PTT, 3.0 g/kg lean mass, 10:1 lactate/pyruvate) performed on 27-28 week old WT and DiC LivKO mice, 4 weeks post-AAV EV/Cre injection, fed NCD (L, N) or WD (M and O) for 21 weeks. Data are presented as mean ± SEM. Statistical analysis of NMR, blood glucose, insulin, and HOMA-IR results performed by two-way ANOVA with Holm-Sidak post-hoc multiple comparison test within 4- and 18-hour fasting groups with significance denoted as $ p<0.05, $$ p<0.01, $$$ p<0.001, $$$ p<0.0001 for main effects and * p<0.05, ** p<0.01, *** p<0.001, **** p<0.0001 for multiple comparisons. Statistical analysis of L/PTT results performed by t-test for each time point denoted as * p<0.05. N = 9-10 for each group.

Next, after the evaluation of metabolic responses to NCD vs WD feeding was completed, mice were allowed to recover from testing, and on week 17 of diet, mice were treated with AAV-Cre and AAV-EV to generate acutely deleted DiC LivKO and WT control mice (Fig 5A). Following knockout of the DiC protein, which requires 3 weeks after AAV administration, tests to examine glucose homeostasis were repeated during weeks 20-23 of diet. DiC LivKO did not alter total or fat mass in NCD or WD mice (Fig 5H). Conversely, DiC LivKO strikingly decreased post-absorptive blood glucose, insulin, and HOMA-IR scores (Fig 5I-K). This was accompanied by decreased GNG in WD-fed mice as reflected by glucose excursion in lactate/pyruvate tolerance tests (10:1 ratio, 3 g/kg lean mass) (Fig 5L-O). These results are consistent with DiC LivKO normalizing blood glucose in WD-fed mice by decreasing GNG, which may in turn decrease insulin production because of lower blood glucose levels.

### DiC LivKO decreases glucose production from ^13^C-lactate/-pyruvate in NCD and WD mice without apparent compensation by the kidney

Given the partially preserved glucose excursion response during L/PTTs in NCD and WD fed DiC LivKO mice (Fig 5L-O), we considered possible mechanisms of gluconeogenic compensation. While the liver is the primary gluconeogenic organ, the kidney also contributes to GNG. To address potential renal compensation and as a concluding experiment after 23 weeks of diet treatment (Figure 5A), we traced ^13^C-lactate/-pyruvate (9:1 ratio, 50% U^13^C labeled, 3 g/kg lean mass) in NCD- and WD-fed mouse livers and kidneys. At 30 minutes post intraperitoneal injection with ^13^C-lactate/-pyruvate, blood glucose and lactate decreased in DiC LivKO mice but did not reach statistical significance in this experiment (Fig 6A-B). Nonetheless, in liver tissue collected at 30 minutes, DiC LivKO decreased glucose abundance and percent ^13^C-enrichment in both NCD and WD fed mice (Fig 6C-F, Table S4). In kidney tissue, percent enrichment of glucose was similarly decreased (Fig 6G-J, Table S4). Given that glucose ^13^C-enrichments in DiC LivKO livers and kidneys decreased similarly, these data suggest that the kidney does not compensate for decreased GNG in liver and that kidney ^13^C glucose enrichments are predominantly downstream of hepatic GNG.

**Figure 6:**
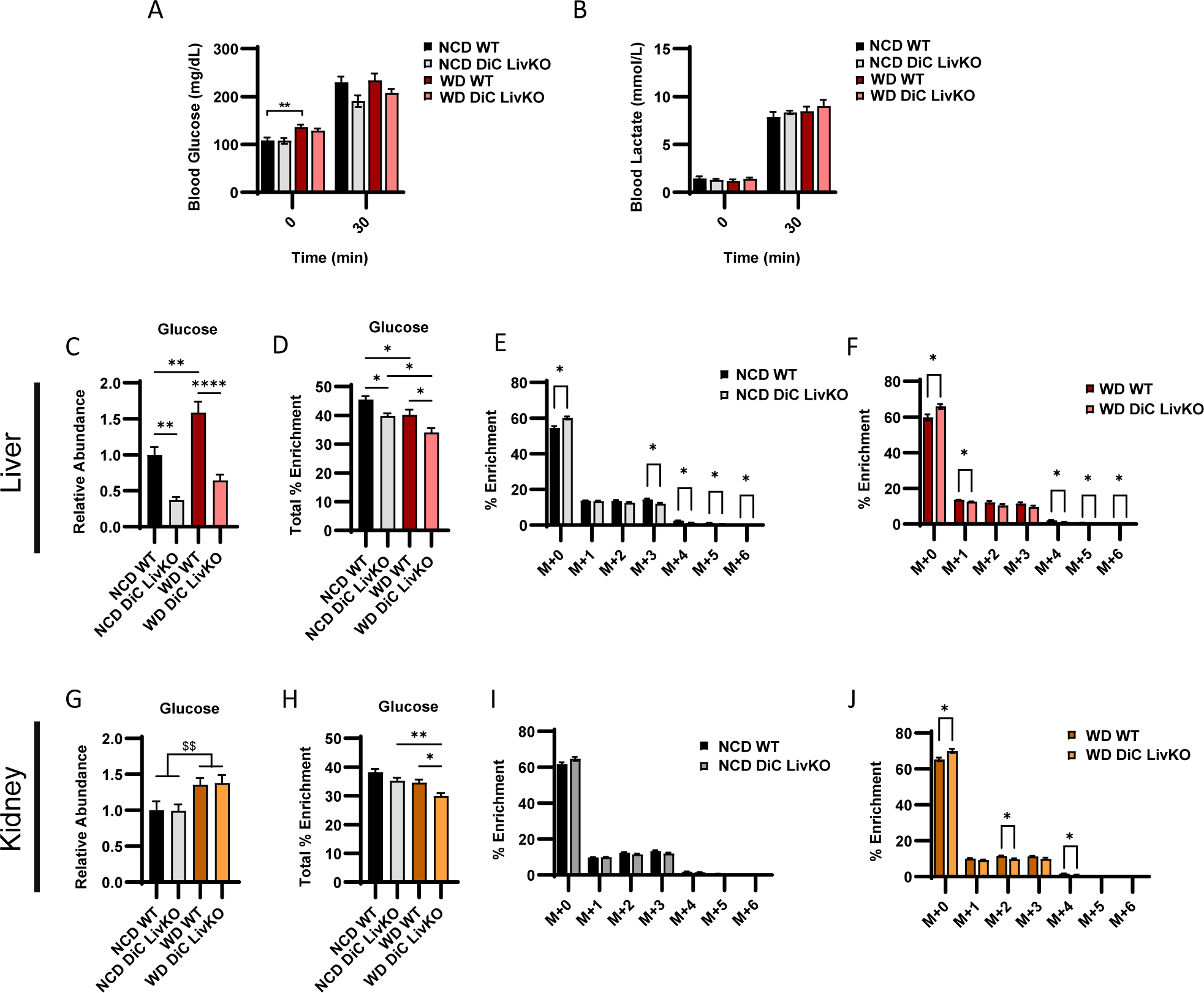
DiC LivKO decreases glucose production from ^13^C-lactate/-pyruvate in NCD and WD fed mice with no apparent compensation by the kidney. 29-30 week old WT and DiC LivKO mice fed normal chow (NCD) or western diet (WD) for 23 weeks were administered ^13^C-lactate/pyruvate tracer (3.0 g/kg lean mass, 9:1 lactate/pyruvate) and euthanized 30 minutes later, 6 weeks post-AAV EV/Cre injection. A-B. Blood glucose (A) and lactate (B) measured by strip meter before tracer administration and at euthanasia. C-J. Terminal liver (C-F) and kidney (G-J) glucose abundance (C and G) total percent ^13^C enrichment (F and H) and mass isotopologue distribution (MID) analysis for NCD (G and I) and WD (H and J) fed groups measured by LC-MS. Data are presented as mean ± SEM. Statistical analysis of blood glucose and lactate, glucose abundance, percent total enrichment by two-way ANOVA with Holm-Sidak post-hoc multiple comparison test within NCD and WD groups with significance denoted as $$ p<0.01 for main effects and * p<0.05, ** p<0.01, **** p<0.0001 for multiple comparisons. Statistical analysis of MIDs by t-test for each time point denoted as * p<0.05. N = 9-10 for each group.

## DISCUSSION

In this study, we show that the mitochondrial dicarboxylate carrier (DiC) mediates in vivo hepatic gluconeogenesis. Previous biochemical studies suggested a significant role for mitochondrial malate export through the DiC in GNG ^16,21^. However, these studies were performed ex vivo, limiting application to normal and T2D physiology. To address this gap, we generated a liver-specific DiC knockout (DiC LivKO) mouse model and tested glucose production from lactate/pyruvate. Hepatic DiC deletion decreased glucose excursion after lactate/pyruvate tolerance tests. This deficit in glucose excursion was rescued by ectopic, in vivo hepatic expression of AAV-delivered WT but not transport null DiC, indicating the transport function of the DiC facilitates its role in GNG. Importantly, we also observed that DiC LivKO decreased flux of ^13^C-lactate/-pyruvate into hepatic and plasma but not kidney glucose, directly demonstrating an in vivo role of the DiC in hepatic GNG.

In addition to demonstrating a role for the DiC in GNG, a key finding of our study is that the DiC contributes to elevated blood glucose and insulin levels induced by high fat, high sucrose Western diet (WD) feeding. Previous work has shown that mitochondrial pyruvate import by the mitochondrial pyruvate carrier (MPC) facilitates GNG from pyruvate ^30,31^. Furthermore, MPC deletion impairs hepatic GNG and improves glucose tolerance in mouse models of T2D ^30,31^. Together with previous findings on the MPC, the results of this investigation show that an MPC-DiC gluconeogenic flux axis mediates pyruvate-driven GNG that contributes to hyperglycemia in T2D states. A notable finding of our current study is that acute genetic deletion of the DiC after the induction of diet-induced obesity, which mimics pharmacologic disruption more accurately than constitutive deletion, corrected hyperglycemia, hyperinsulinemia, and HOMA-IR scores in WD-fed mice.

Although much more work will be required to understand if the DiC is a viable therapeutic target, several properties of the DiC align with desirable features of a target to modulate GNG. In humans, DiC protein abundance is greatest in the liver, followed by the kidney and duodenum, the major gluconeogenic organs ^32^. Thus, DiC inhibition could exert the greatest biological effect in its primary target organ, the liver. Moreover, deleting the DiC in the liver does not alter baseline fasting blood glucose levels. This suggests that DiC inhibition suppresses excess but not homeostatic glucose output in T2D, an ideal characteristic of a therapeutic target to decrease GNG.

Our study has advantages and limitations. An advantage is that compared to previous biochemical studies using non-specific inhibitors in cell culture and ex vivo tissue systems, the in vivo, hepatocyte-delimited deletion employed here provides strong evidence that the DiC mediates hepatic GNG in vivo. This conclusion is further validated by the in vivo DiC LivKO rescue experiments we performed with ectopic hepatic expression of DiC WT and null constructs, which showed nearly complete and no rescue, respectively, of GNG and related TCA cycle metabolic features. A limitation of this study is that our ^13^C tracing results show that DiC deletion partially decreases but does not block pyruvate-driven GNG, which demonstrates the existence of an alternative mitochondrial carbon export pathway(s) to supply GNG. Whether and how the DiC and other export pathways supplying GNG are discretely regulated will need to be resolved to fully understand the roles of each. Overall, while outstanding questions remain, the results of this study provide greater understanding of carbon trafficking mechanisms supplying GNG in normal and T2D states.

## ACKNOWLEDGEMENTS

We are grateful to the University of Iowa Fraternal Order of Eagles Diabetes Research Center Metabolomics Core Facility for technical assistance. This work was supported by grants NIH R01 DK104998 (EBT), NIH R01 DK138664 (EBT), University of Iowa Healthcare Distinguished Scholar Award (EBT), NIH R01 DK126817 (LY), AHA CDA851976 (AJR), T32 GM007337 to Steven Lentz and AHA 23PRE1020471 (DJP), NIH T32 DK112751 to Andrew Norris (KCHF), and P30CA086862 to Mark Burkard, which contributed to support of core facilities utilized for this research.

## AUTHOR CONTRIBUTIONS

EBT conceived the project. DJP, KCFH, LY, and EBT designed experiments. DJP, KCFH, RAM, AJR, AA, QQ, GRM, and PS performed experiments. DJP, KCFH, RAM, AJR, AA, and EBT performed data analysis and data visualization. DJP, KCFH, and EBT wrote the manuscript. DJP, KCFH, AJR, and EBT edited the manuscript.

## RESOURCE AVAILABILITY

### Lead contact

Additional information and requests for reagents and resources should be directed to and will be fulfilled by the lead contact, Dr. Eric Taylor.

### Materials availability

New reagents generated in this study are available by request to the lead contact.

## MATERIALS AND METHODS

### Animal Studies

All animal experiments were approved by the University of Iowa Office of the Institutional Animal Care and Use Committee (IACUC). Mice were fed normal chow diet (NCD) (Teklad, 2920x), or, when indicated, a high-fat, high-sucrose Western Diet (WD) (Research Diets, D12079B). C57Bl/6J mice were purchased from Jackson Laboratories.

### Generation of mouse lines

Cryopreserved *Slc25a10^tm1a(EUCOMM)Wtsi^* sperm were obtained from the European Conditional Mouse Mutagenesis Program (EUCOMM). Oocytes were isolated from C67BL/6J mice, and in vitro fertilization was performed by the Genome Editing Facility at the University of Iowa. Two-cell stage embryos were implanted into recipient females for gestation. The resultant heterozygous *Slc25a10^tm1a(EUCOMM)Wtsi/+^* knockout-first mice were bred to mice expressing Flp recombinase under the ROSA promoter generating *Slc25a10^tm1c/+^* conditional allele mice. *Slc25a10^tm1c/+^* mice were again bred to C57Bl/6J mice to outcross the Flp transgene, and the resulting heterozygous *Slc25a10^tm1c/+^*were crossed to generate homozygous *Slc25a10^tm1c/tm1c^* (*DiC^fl/fl^*) mice used in this study. To study whole-body DiC disruption, knockout-first allele *Slc25a10^tm1a(EUCOMM)Wtsi/+^* were intercrossed generating *Slc25a10^+/+^*(WT), *Slc25a10^tm1a(EUCOMM)Wtsi/+^* (KO1/WT), and *Slc25a10^tm1a(EUCOMM)Wtsi /tm1a(EUCOMM)Wtsi^* (KO1/KO1).

### Generation of DiC LivKO mice

Liver specific DiC knockout mice (DiC LivKO) and littermate paired control (WT) mice were generated as previously reported ^30^. *DiC^fl/fl^*mice were anesthetized and a received retroorbital injection of AAV8-TBG-Cre to hepatically express Cre recombinase or empty vector AAV8-TBG-NULL. For most experiments, virus was delivered at 1*10^11^ genome copies (GC) diluted in 100 µL PBS per mouse. To generate DiC LivKO subsequent to WD feeding, 2*10^11^ GC of virus was diluted in 100 µL PBS per mouse.

### DiC Expression Vector Construction

The cDNA sequence encoding the major DiC splice isoform (NM_012140.5, DiC-WT) was cloned into an AAV vector and expressed via the liver-specific TBG promoter. For transport null DiC (DiC-R261Q), a R261Q mutation was introduced into the DiC-WT cDNA sequence by altering the CGT arginine encoding codon to CAG, which encodes glutamine. This mutation is orthologous to the *Slc25a15* (Orc1) R275Q mutation known to abolish transport activity without disrupting protein abundance ^25–27^.

### DiC In Vivo Expression

For in vivo experiments, *DiC^fl/fl^* mice received a single retro-orbital injection containing two constructs: 1) AAV8-TBG-Cre or AAV8-TBG-NULL to delete endogenous DiC and 2) DiC-WT, DiC-R261Q, or empty vector (EV). 1*10^11^ GC of each construct was injected for a total of 2*10^11^ GC diluted in 100 µL PBS per mouse.

### Body Composition by NMR

Lean mass and fat mass were measured using the LF50 Bruker MiniSpec housed in the University of Iowa Metabolic Phenotyping Core.

### Lactate/Pyruvate Tolerance Test

At ZT12 the day prior to the tolerance test, mice were single-housed and food was removed, with free access to water. The next day, a 10:1 (w/w) lactate/pyruvate mixture was prepared in sterile water. This solution was pH adjusted to 7.4 and filter sterilized. At ZT5, mice were brought to the procedure room and weighed. At ZT6, blood was sampled from tail vein to measure glucose and lactate. Blood glucose was measured with a handheld glucometer (OneTouch UltraMini) and glucose testing strips (OneTouch Glucose Strips). Blood lactate was measured with a handheld lactate meter (Lactate Plus, #40828) and testing strips (Lactate Plus, #40813). Following these measurements, mice were intraperitoneally injected with a lactate/pyruvate solution (10:1 or 9:1 ratio to approximate endogenous redox balance) at a dosage of 2.2 g/kg body weight (based on weights at ZT5) or 3.0 g/kg lean mass based on NMR body composition measurements a few days prior (dosing is indicated in figure legends). Blood glucose and lactate measurements were repeated at 15, 30, 60, 90, and 120 minutes following lactate/pyruvate administration.

### Glucose Tolerance Test

At ZT3 the day of the tolerance test, mice were single-housed and food was removed. A 10% (w/v) glucose solution was prepared in sterile water and filter sterilized. At ZT6, mice were brought to the procedure room and weighed. At ZT7, blood glucose measurements were taken as described in the lactate/pyruvate tolerance test. Mice were then intraperitoneally injected with glucose at a dosage of 1.25 g/kg body weight (based on weights taken at ZT6). Following glucose administration, blood glucose measurements were repeated at 15, 30, 45, 60, 90, and 120 minutes.

### Insulin Tolerance Test

At ZT3 the day of the tolerance test, mice were single-housed and food was removed. 0.1 U/mL of Humulin R was prepared in 0.9% sterile saline. At ZT6, mice were brought to the procedure room and weighed. At ZT7, blood glucose measurements were taken as described in the lactate/pyruvate tolerance test. Mice were then intraperitoneally injected with insulin at a dose of 0.75 U/kg body weight (based on weights taken at ZT6). Following insulin administration, blood glucose measurements were repeated at 15, 30, 45, 60, 90, and 120 minutes.

### Glucose Homeostasis Measurements

At ZT12 the day prior to the assay (18 hour fasted measurements) or ZT2 the day of the assay (4 hour fasted measurements), mice were single-housed and food was removed. At ZT5, mice were brought to the procedure room. Plasma collection began at ZT6. Blood glucose was measured as previously described for the lactate/pyruvate tolerance test. The tail vein was then sampled to collect whole blood into EDTA coated tubes. Blood samples were centrifuged for 3000 x g for 15 minutes at 4°C. The plasma compartment in the resulting supernatant was collected and frozen at −80°C. The CrystalChem Ultra Sensitive Mouse Insulin ELISA kit was used to measure insulin. Specifically, 4 μL of plasma from each sample was used for each technical duplicate. The homeostatic model assessment for insulin resistance (HOMA-IR) score was calculated by the following formula: HOMA-IR = (Blood Glucose (mmol/L)*Plasma Insulin (µIU/mL))/22.5.

### ^13^C-Lactate/Pyruvate Tracing

At ZT12 the day prior to the tolerance test, mice were single-housed and food was removed, with free access to water. The next day, a 9:1 (w/v) lactate/pyruvate mixture with 50% U-^13^C was prepared by combining equal volumes of 100% ^12^C and 100% U-^13^C lactate/pyruvate solutions. This solution was pH adjusted with HCl to achieve a final pH of 7.4 and filter sterilized. At ZT5, mice were brought to the procedure room and weighed. At ZT6, tail vein blood was sampled to measure glucose and lactate followed by intraperitoneal injecting mice as previously described for the lactate/pyruvate tolerance test. At 27 minutes post-injection, mice were anesthetized with isoflurane and blood glucose and lactate measurements were repeated. 30 minutes post-injection, the left lateral lobe of the liver was dissected with diaphragm intact and immediately (within 2 seconds) frozen using liquid nitrogen cooled freeze clamps as previously described ^33^. For experiments involving kidney harvest for ^13^C tracing, a kidney was dissected about 10 seconds after liver tissue and frozen immediately with liquid nitrogen cooled freeze clamps. Blood was obtained via cardiac puncture and placed into tubes with 10 µL 0.5 M EDTA, pH 8.0. Blood samples were centrifuged for 3000 x g for 15 minutes at 4°C. The plasma compartment in the resulting supernatant was collected and frozen at −80°C.

### Steady-State Metabolomic Profiling

At ZT0 on the day of tissue harvest, mice were single-housed and food was removed. At ZT3, mice were transferred to the procedure room. The experiment began at ZT4. Liver harvest and blood collection were performed as previously described for the ^13^C-lactate/pyruvate tracing study.

### Metabolite Extraction from Liver and Kidney Tissue

Frozen liver or kidney tissue (30-80 mg) was weighed without thawing and placed into 1.7 mL microcentrifuge tubes with needle punctured caps. To remove water, these samples were then lyophilized overnight at −80°C and with a pressure of less than 0.3 torr. A 2:2:1 methanol:acetonitrile:water extraction buffer was then added to lyophilized tissue at a volume in microliters 18 times the sample mass in milligrams. Samples were then homogenized using a BeadRupter bead mill homogenizer (Omni International) for 30 seconds at 6.45 MHz. Samples were rotated for 1 hour at −20°C. Afterwards, samples were centrifuged at 21,100 x g at 4°C for 10 minutes to pellet debris. The resulting supernatant was transferred to new 1.7 mL microcentrifuge tubes and centrifuged again for 21,100 x g at 4°C for 10 minutes to pellet any remaining debris. The supernatant was transferred to new tubes and vortexed. For LC-MS, 300 μL of this extract was placed in a new 1.7 microcentrifuge tube and dried until all solvent evaporated (∼2 hours) using a Speedvac Vacuum concentrator (Thermo) without heating. Pooled quality control (QC) sample was prepared by combining an equal volume of each extract, which was dried similar to the other samples.

### LC-MS analysis

After drying, extracts were resuspended in 30 μL of a 1:1 acetonitrile:water mixture and vortexed for 10 minutes. Extracts were then incubated at −20°C overnight. The next day, samples were centrifuged at 21,100 x g at 4°C for 15 minutes. The resulting supernatant was transferred to autosampler vials for LC-MS analysis. For each prepared sample, 3-4 µL was separated using a Millipore SeQuant ZIC-pHILIC (2.1 × 150 mm, 5 µm particle size, Millipore Sigma #150460) column with a ZIC-pHILIC guard column (20 x 2.1 mm, Millipore Sigma #150437) attached to a Thermo Vanquish Flex UHPLC. The mobile phase comprised Buffer A [20 mM (NH4)2CO3, 0.1% NH4OH (v/v)] and Buffer B [acetonitrile]. The chromatographic gradient was run at a flow rate of 0.150 mL/min as follows: 0–21 min-linear gradient from 80 to 20% Buffer B; 21-21.5 min-linear gradient from 20 to 80% Buffer B; and 21.5–28 min-hold at 80% Buffer B. Data was acquired using a Thermo Q Exactive MS with a spray voltage set to 3.0 kV, the heated capillary held at 275°C, and the HESI probe held at 350°C. The sheath gas flow was set to 40 units, the auxiliary gas flow was set to 15 units, and the sweep gas flow was set to 1 unit. MS data resolution was set at 70,000, the AGC target at 10e6, and the maximum injection time at 200 ms. For metabolite profiling, the mass spectrometry operated in Full Scan mode with polarity switching. For ^13^C tracing experiments, the mass spectrometry operated in targeted selected ion monitoring (tSIM) mode either with separate negative or positive polarity or with polarity switching. To measure instrument drift, the same QC sample was analyzed at the beginning, equally throughout, and at the end of the run.

### Mass Spectrometry Data Analysis

MS data were processed using the Thermo Scientific TraceFinder (5.1) software. Targeted metabolites were identified based on the University of Iowa Metabolomics Core facility standard-confirmed, in-house library defining a target ion accurate mass, retention time, and MS/MS fragmentation pattern. ^13^C isotopologues were identified as M+1.0033 m/z per carbon labeled ions of standard-confirmed metabolites. The NOREVA tool was used to correct for instrument drift by using the QC sample analyzed throughout the instrument run ^34,35^. For ^13^C-tracing analysis in non-diet cohorts, ^13^C-natural abundance was corrected using liver samples from non-tracer injected mice analyzed simultaneously with tracer injected liver samples. For ^13^C-tracing analysis in Western Diet mice, ^13^C-natural abundance was corrected using Isocor 2.2.1 ^36,37^. The following settings were used: Isotope tracer: ^13^C, high resolution. High resolution parameters: Resolution formula: orbitrap, instrument resolution: 70000, at m/z: 400.0. There was no correction for natural abundance of the tracer element. Isotopic purity of the tracer: ^12^C: 0, ^13^C: 1.

### Mitochondrial Isolation and Protein Extraction

Mitochondria were isolated from frozen liver and kidney tissue in an isolation buffer (MIB) comprising 70 mM sucrose, 210 mM D-mannitol, 5 mM HEPES, 1 mM EGTA, and 0.1% bovine serum albumin (BSA) in sterile water and pH adjusted to 7.4 with KOH. MIB was supplemented with 1 mM dithiothreitol (DTT) and 1x ProteaseArrest protease inhibitor cocktail. MIB was added to frozen liver tissue at a volume (microliters) 40 times tissue weight in milligrams and immediately homogenized. The resulting homogenates were then centrifuged at 800 x g for 10 minutes at 4°C. The supernatant was then transferred to a new tube and centrifuged for 8000 x g for 10 minutes at 4°C. Following this step, the supernatant was removed, and the pellet was resuspended in MIB at the same volume used for homogenization. This resuspension was then centrifuged at 8500 x g for 10 minutes at 4°C. After removing the supernatant, the pellet was resuspended in MIB at the same volume used for homogenization. This resuspension was centrifuged at 9000 x g for 10 minutes at 4°C. The supernatant was then aspirated, and the pellet was frozen at - 80°C. For protein extraction, the mitochondrial pellet was resuspended in 30 μL of RIPA lysis buffer and rotated for 30 minutes at 4°C. Samples were then centrifuged at 21,100 x g for 10 minutes at 4°C. The supernatant was recovered and used for analysis. Protein concentration was quantified using Bradford reagent (BioRad, 5000006) measured at 595 nm. The resulting lysates were then used for western blotting.

### Western Blotting

12% Tris-Glycine SDS-PAGE gels with a 5% stacking gel were prepared and stored at 4°C at least one day prior to western blotting. Protein lysates were diluted in water and combined with 4x Laemmli buffer containing beta-mercaptoethanol. Lysates were then heated for 10 minutes at 50°C and loaded onto gels. Proteins were separated by running samples for 30 minutes at 50 V followed by 2 hours at 100 V. Proteins were transferred to a 0.45 µm nitrocellulose membranes (Amersham, 10600002), blocked with a 1% nonfat dry milk/1% bovine serum albumin (BSA) solution in TBST (1 mM Tris, 150 mM NaCl, 0.1% Tween-20) and then probed overnight with primary antibody. The next day, membranes were washed with TBST and incubated with fluorescent secondary antibodies for 1 hour at room temperature. Proteins were imaged using the Li-Cor Odyssey CLx system and densitometry was performed using ImageJ.

### Statistical analysis

Data were then analyzed by unpaired t-tests, one- or two-way ANOVA with the Holm-Sidak post-hoc multiple comparison test applied as needed for hypothesis testing. Statistically significance differences were defined as having p-values < 0.05 (^*,$^), p < 0.01 ^(**, $$^) and p < 0.001 ^(***,$$$^). Results were plotted as mean ± standard error of the means (SEM). The number of biologically unique samples and statistical test used in each experiment is specified in the relevant figure legend.

